# Probabilistic Thermodynamic Analysis of Metabolic Networks

**DOI:** 10.1101/2020.08.14.250845

**Authors:** Mattia G. Gollub, Hans-Michael Kaltenbach, Jörg Stelling

**Affiliations:** Department of Biosystems Science and Engineering and SIB Swiss Institute of Bioinformatics, ETH Zurich, Basel, 4058, Switzerland

## Abstract

Random sampling of metabolic fluxes can provide an unbiased description of the capabilities of a metabolic network. However, current sampling approaches do not model thermodynamics explicitly, leading to inaccurate predictions of an organism’s potential or actual metabolic operations. We present a probabilistic framework combining thermodynamic quantities with steady-state flux constraints to analyze the properties of a metabolic network. It includes methods for probabilistic metabolic optimization and for joint sampling of thermodynamic and flux spaces. Applied to a model of *E. coli*, we use the methods to reveal known and novel mechanisms of substrate channeling, and to accurately predict reaction directions and metabolite concentrations. Interestingly, predicted flux distributions are multimodal, leading to discrete hypotheses on *E. coli* ‘s metabolic capabilities. C++ source code with MATLAB interface available at https://gitlab.com/csb.ethz/pta.

## 1 Introduction

Constraint Based Models (CBMs) aim to characterize the metabolic capabilities of an organism, potentially at genome-scale, by predicting metabolic fluxes based on constraints arising from the metabolic network’s structure [1]. Because many degrees of freedom exist for the network’s (potential, if not practical) operation, a central object of the analysis is the flux space. Formally, the flux space *ℱ* of a metabolic network with *n* reactions and *m* metabolites is the set of flux distributions **v** ∈ ℝ^*n*^ that satisfy steady state and capacity constraints:

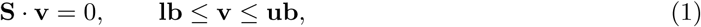

where **S** ∈ ℝ^*m×n*^ is the stoichiometric matrix and **lb** ∈ℝ^*n*^ and **ub** ∈ℝ^*n*^ are measured or assumed lower and upper bounds on fluxes.

The most common approaches to analyzing *ℱ* predict particular flux solutions, for example, by assuming cellular objectives in Flux Balance Analysis (FBA) [1]. To explore and characterize all possible solutions that lead to observed phenotypes or desired goals, Uniform Sampling (US) [2, 3] and its extensions in the form of loopless [4, 5] and non-uniform sampling [6–8] provide an unbiased description of the flux space.

However, the network structure alone often constrains the flux space insufficently. Different - omics data from high-throughput measurement technologies can in principle be used to derive additional, condition-specific constraints, but their integration is challenging [9]. In contrast, constraints based on thermodynamics derive from first principles; they are absolute and independent of specific kinetic properties or regulatory mechanisms. Combined with the network structure, modeling thermodynamic laws has the potential to reveal the options available to the cell.

In a metabolic model, thermodynamics quantifies the favourability of a set Γ of *γ* reactions (excluding unbalanced pseudo-reactions such as biomass and exchange reactions) with Gibbs reaction energies **Δ**_**r**_**G**′ ∈ℝ^*γ*^ :

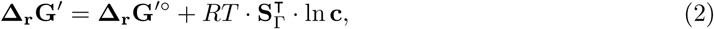

where **c** ∈ℝ^*m*^ are the metabolite concentrations, **Δ**_**r**_**G**^′°^ ∈ℝ^*γ*^ the standard Gibbs reaction energies (corrected for pH, electrostatic potential, and ionic strength), *R* the gas constant, *T* the temperature, and 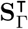 is the transpose of the stoichiometric sub-matrix corresponding to Γ. The second term of Eq. (2) (here called the *activity term*) maps the contribution of metabolite concentrations to the reactions they participate in. Reaction energies constrain fluxes via the second law of thermodynamics, which implies that positive (negative) net flux can only occur if the free energy of a reaction is negative (positive). This ensures that biochemical processes result in net energy dissipation.

How well thermodynamics constrain a model depends on the precision of the estimates of the variables in Eq. (2). Standard reaction energies can be estimated from measured equilibrium constants [10] by the group contribution method and its extensions [11–13]. Because multiple reactions may affect the same chemical groups, their estimates are often linearly dependent and thus lie in a lower-dimensional subspace (Fig. 1A). Metabolomics data can provide precise estimates or approximate distributions [14] for metabolite concentrations (Fig. 1B).

**Figure 1:**
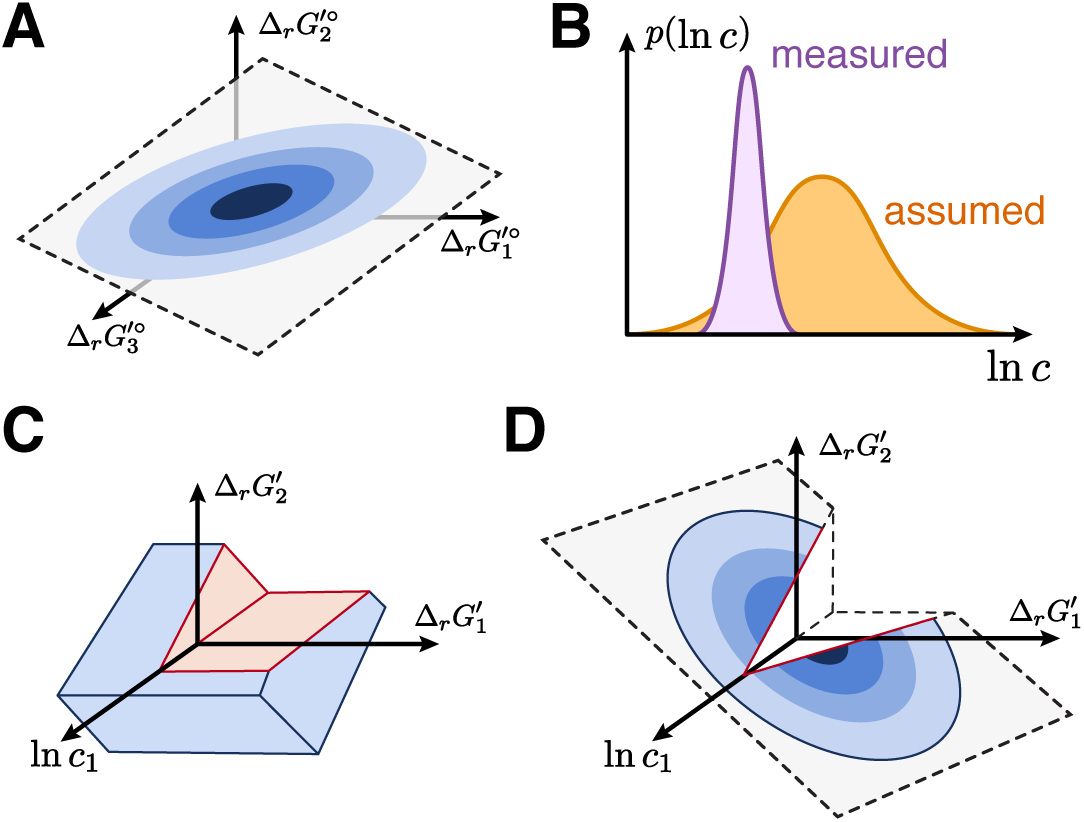
Integration of thermodynamics in CBMs. (**A**) Group contribution methods provide estimates of standard reaction energies. The uncertainties of multiple estimates may be correlated and lie in a lower dimensional subspace. (**B**) Probability distributions of metabolite concentrations can be constructed from measured or assumed data. (**C**) The space of reaction energies and metabolite concentrations defined by TMFA, with regions that are not feasible at steady state removed (red). The space is defined by independent bounds on each variable of ln **c** and **Δ**_**r**_**G**^′°^, leading to an overapproximation of the uncertainty. (**D**) PTA models the complete uncertainty information probabilistically, leading to a joint probability distribution over the reaction energies of the entire network. Steady state flux constraints restrict it to feasible orthants.

Thermodynamics-based Metabolic Flux Analysis (TMFA) [15] integrates thermodynamics into CBMs with (i) a set of linear constraints describing Eq. (2) and bounds on its quantities and (ii) integer constraints enforcing the second law of thermodynamics. TMFA has been used to predict metabolite concentrations [14, 15] and pathway favourability [16, 17]. However, it describes the uncertainty by independent error bounds for each variable. This facilitates the construction of optimization problems, but also over-approximates the uncertainty. Fig. 1C illustrates a space of thermodynamic variables in TMFA: independent constraints on ln **c** and **Δ**_**r**_**G**^′°^ constrain **Δ**_**r**_**G**′ as well. In addition, the steady state condition (Eq. (1)) excludes orthants that imply directions without a feasible flux distribution in *ℱ*. This approach discards any correlation between the 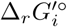 of different reactions and the probability of a particular solution.

The common construction of CMBs for pure flux-based analysis poses a further challenge. Model curators or automatic pipelines regularly need to prevent unrealistic internal or ATP-generating cycles [18]. Because the flux balance framework does not resolve this problem consistently, assumptions or condition-specific biological knowledge are employed to assign reaction reversibilities. Importantly, missing knowledge or inaccurate irreversibilities can impose false thermodynamic constraints that do not affect flux balance and thus remain undetected.

To address these challenges, here we propose *Probabilistic Thermodynamic Analysis* (PTA), a framework in which the uncertainty of free energies and concentrations is modeled with a joint probability distribution (Fig. 1D). We develop optimization and sampling methods based on PTA and show their application to predict substrate channeling, reaction directions, and metabolite concentrations, as well as to explore the capabilities of the metabolic network.

## 2 Methods

### 2.1 The steady-state thermodynamic space

To account for correlations between the uncertainties of reaction energies of different reactions, we assume that metabolite concentrations follow a log-normal distribution based on measurements or an assumed physiological distribution [14]

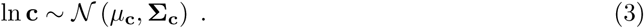

From the estimates of a group contribution method [12], we obtain a normal distribution for the standard reaction energies

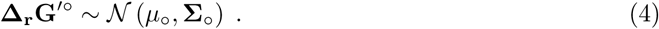

From Eq. (2), it follows that the reaction energy estimates are also normally distributed as

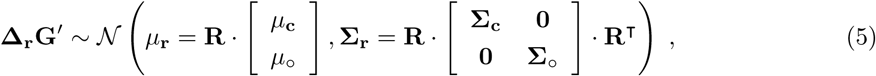

where 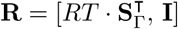. For computational convenience, we define the vector **t** = [ln **c, Δ**_**r**_**G**^′°^, **Δ**_**r**_**G**′]^**T**^. It is again normally distributed with mean *μ*_**t**_ = [*μ*_**c**_, *μ*_°_, *μ*_**r**_]^**T**^ and covariance matrix

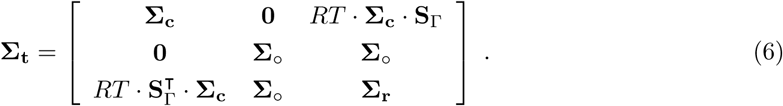

We use this distribution to constrain **t** to values that are compatible with the estimated concentrations and standard free energies up to a confidence level *α*, such that

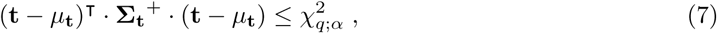

where **Σ**_**t**_^+^ is a pseudo-inverse of **Σ**_**t**_, of rank *q* = rank(**Σ**_**t**_^+^) = rank(**Σ**_**t**_), and 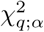 is the *α*-quantile of the *χ*^2^-distribution with *q* degrees of freedom. We exploit that *q* is much smaller than the dimension of **t** by using a *q*-dimensional standard normal distribution to express **t** as

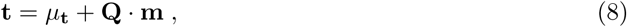

where **Q** = **Σ**_**t**_^1/2^ is a matrix square root such that **Σ**_**t**_ = **QQ**^**T**^.

Lastly, we enforce the second law of thermodynamics and require that all reactions in Γ have a well-defined direction, that is, non-zero flux. For each *i*, 1 *≤ i ≤ γ*:

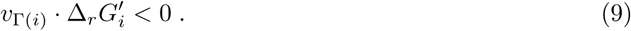

Note that this requires blocked reactions to be removed in advance.

We define the *steady-state thermodynamic space* (or short: *thermodynamic space*) *𝒯* as the set of **t** that fulfill Eq. (7) and satisfy the steady state constraints Eq. (1), Eq. (9). In other words, *𝒯* defines the possible reaction energies and metabolite concentrations at steady-state.

### 2.2 Probabilistic Metabolic Optimization (PMO)

The definitions of *ℱ* and *𝒯* allow us to extend the FBA framework to include thermodynamic quantities similar to TMFA. We translate Eq. (1) and Eq. (7) into linear and quadratic constraints and rewrite Eq. (9) with integer constraints in the *big-M* formulation [19]. The resulting Mixed-Integer Quadratically-Constrained Program (MIQCP) can be used to perform FBA-like analyses under thermodynamic constraints.

Since we know the probability distribution of **t**, we can also determine the most probable reaction energies and metabolite concentrations by setting a quadratic objective that maximizes the probability of **t** under steady-state constraints:

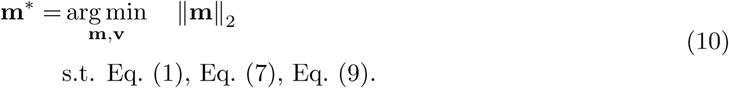

We call **t**^*∗*^ = *μ*_**t**_ + **Q** · **m**^*∗*^ the *mode* of the distribution of **t**.

### 2.3 Thermodynamic assessment of metabolic models

*Structural assessment* aims to detect *thermodynamic inconsistencies* in a model, that is, groups of reactions that, given their irreversibility annotation, cannot satisfy thermodynamic constraints with any assignment of concentrations and standard free energies. For example, two stoichiometrically equivalent paths can be irreversible in opposite directions; then, all non-zero flux solutions contain a thermodynamically infeasible internal cycle. To resolve such cases, we computed an Irreducible Inconsistent Subsystem (IIS) of the optimization problem, i.e., a minimal set of constraints such that, if any constraint is removed, the problem becomes feasible. We then inspected the reactions in the IIS to find and resolve incompatible irreversibilities. If the IIS was too large, we searched for reactions that could participate to internal cycles by setting all exchange reactions to zero and selecting the reactions that could still have non-zero flux. We then manually searched for groups of reactions for which all non-zero flux solutions contained internal cycles and resolved conflicts.

*Quantitative assessment* of a model aims to find features in PMO predictions indicative of knowledge gaps or model inaccuracies. First, we construct a condition-specific model with available flux and concentration constraints and use PMO to estimate **t**^*∗*^. Next, for each predicted value 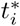 we compute its *z-score z*_*i*_ which measures the deviation between 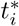 and *μ*_*i*_ in units of standard deviation (see Supplementary Information). We focused on metabolite concentrations and classified predictions with |*z*_*i*_| greater than a threshold *θ* (or *c*_*i*_ *≥* 10*mM* for non-intracellular metabolites) as *anomalies* and investigated whether those indicate missing or unknown mechanisms or model inaccuracies. Finally, we curated the model with the results of the search. The values in **t**^*∗*^ represent the unlikely case in which predicted standard free energies are as close as possible to the mean estimate and reactions can operate close to the thermodynamic equilibrium. Because the predicted deviations likely underestimate the true deviations, we use a conservative *θ* = 1.

### 2.4 Thermodynamics and Flux Sampling (TFS)

We use sampling to characterize *𝒯* in terms of probability distributions of metabolite concentrations as well as probabilities for the different flux modes. Geometrically, *𝒯* is the intersection between an ellipsoid and the orthants with at least one steady-state flux distribution satisfying flux and thermodynamic constraints. This space cannot be defined explicitly, and is usually neither convex nor connected. In fact, an *n*-dimensional space has 2^*n*^ orthants, each corresponding to a possible combination of reaction directions in a metabolic network. Clearly, the enumeration—let alone verification of the steady state condition—of all orthants of *𝒯* is infeasible with many reversible reactions.

We therefore developed a Markov Chain Monte Carlo (MCMC) method based on the Hit-and-Run sampler for General Target Distributions [20] to sample *𝒯* (see algorithm 1).

#### Choice of initial points

For MCMC simulations it is generally advised to simulate multiple chains, choosing the starting points from an over-dispersed distribution, to detect if the chains explore the entire space. For each reversible reaction, we use PMO to find two sets of reaction directions over the network: one that maximizes the flux through the reaction, and one that minimizes it. Then, again using PMO, for each set of directions we search for a point **s**_**i**_ in *𝒯* that satisfies the directionality constraints. We require each **s**_**i**_ to have maximal distance from the boundary of the respective space to reduce the risk of numerical errors. Finally, we start simulating one chain from each **s**_**i**_.

#### Finding the intersection *I*

At each step of Hit-and-Run we first intersect the ray with the ellipsoid defined by Eq. (7) and a set of linear inequalities constraining the signs of free energies of irreversible reactions. Without a close-form description of the boundary of the space, finding the orthants that allow steady-state flux solutions is challenging. Despite the exponential number of orthants, a ray in an *n*-dimensional space only crosses at most *n* + 1 of them. Thus, at each step we identify all the orthants intersected by the ray and discard those that do not allow a steady-state flux distribution. We verify the steady-state condition with a linear program that checks the existence of a flux distribution satisfying Eq. (1) and the directionality constraints of the orthant. For each feasible orthant, we add its intersection with the ray to *I*.

#### Sampling from *I*

We sample the next point along a ray proportionally to the target probability distribution, which is a Multivariate Normal (MVN). First we restrict the MVN to a normal distribution along the 1-dimensional ray. We then compute the Cumulative Distribution Functions (CDFs) for each valid segment of *I* to determine its weight, sample a valid segment according to these weights and finally sample a point in the segments according to the associated truncated normal distribution on the segment. We optionally store the sample and the samples count for the selected orthant to estimate the orthant’s probability.

##### Algorithm 1: Sampling the steady state thermodynamic space *𝒯*

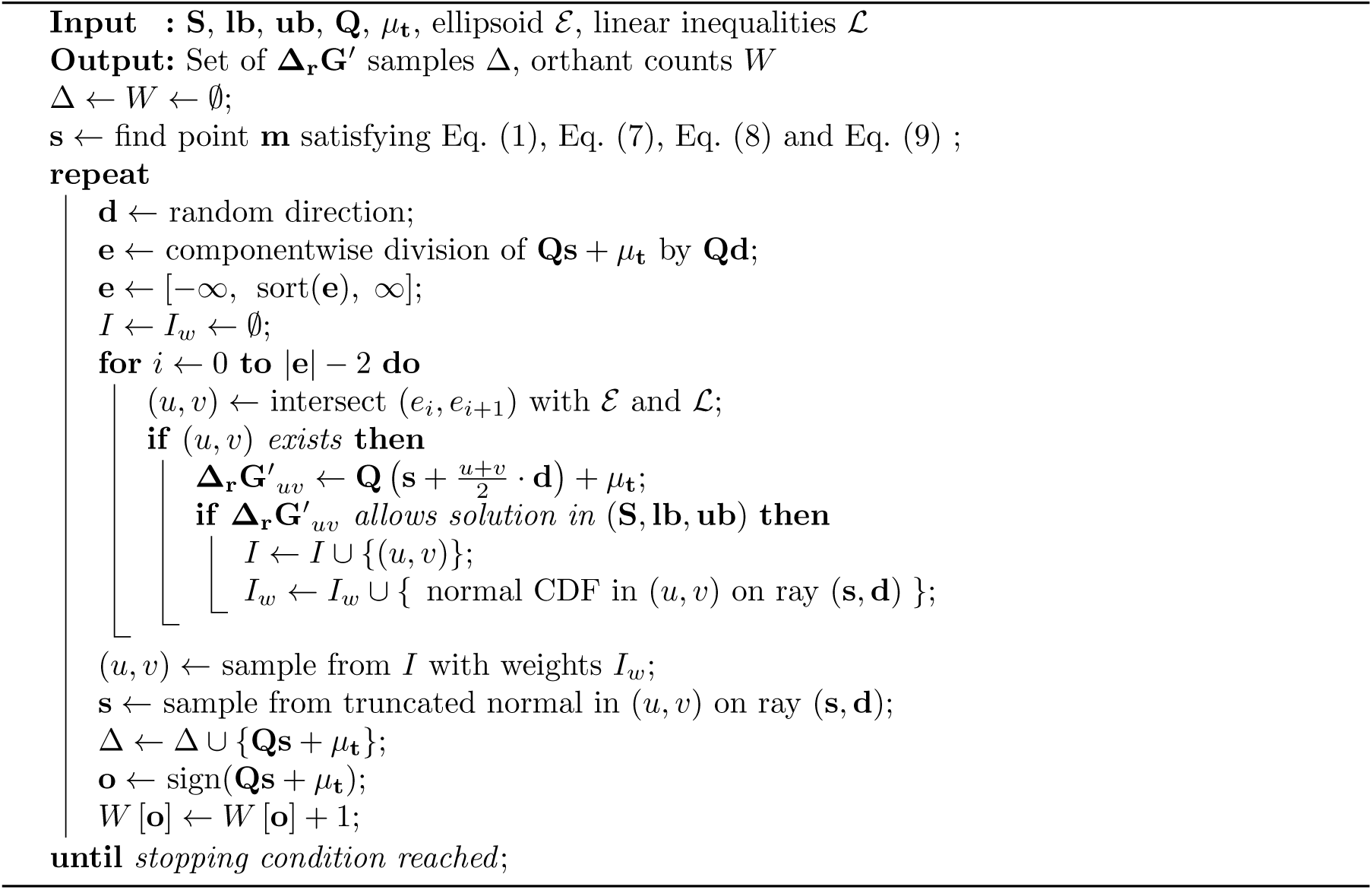

#### Reducing the dimensionality

Our method for sampling *𝒯* so far neglects that only reaction free energies determine flux directions, and that free energies are determined only by concentration ratios, not absolute concentrations. Thus, we can reduce the dimensionality of the problem by not representing metabolite concentrations explicitly and solving Eq. (4) using *μ*_**r**_ and **Σ**_**r**_ instead of *μ*_**t**_ and **Σ**_**t**_. This allows faster convergence of the chains. Afterwards, for each sample of **Δ**_**r**_**G**′_*i*_, we sample the corresponding ln **c** and **Δ**_**r**_**G**^′°^ from the distribution of **t** conditioned on **Δ**_**r**_**G**′_*i*_. This distribution can be constructed from the *Schur’s complement* of **Σ**_**r**_ in **Σ**_**t**_ and has the form of a MVN, which can be sampled efficiently:

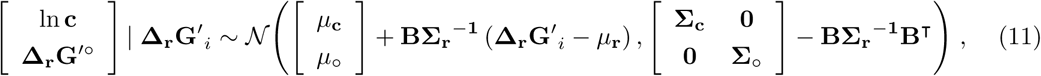

where B =[*RT***Σ**_c_S_Γ_ **Σ**_°_]^T^ is a submatrix of **Σ** _t_.

#### Convergence

Since each sampled orthant is full-dimensional and all orthants are reachable, asymptotic convergence to arbitrary target distributions is guaranteed [20]. However, there is no theoretical bound on the convergence rate. We therefore asses the quality of the samples chains using recommended criteria on the chains’ Potential Scale Reduction Factor (PSRF) and Effective Sample Size (ESS) [21]. Several hit-and-run samplers reformulate the problem in isotropic parametrization to improve convergence [3]. Since the distribution of **m** is a standard MVN, the parametrization already has a potentially high isotropy but this may not be optimal because flux constraints exclude parts of the solution space. To improve convergence, we initially run shorter chains and compute the PSRF of each reaction energy. We then adjust the parametrization such that the space is better explored along dimensions showing high PSRFs.

#### Flux sampling with thermodynamic prior

Finally, we sample the flux space using the estimated distributions of reaction directions as prior. We assume that the probability of an orthant is determined by its probability in the thermodynamic space, for which we already obtained estimates. For each direction sample **d**_**i**_ ∈ *{*−1, 1*]γ*, one can easily construct and sample the flux space defined by Eq. (1) and constrained by **d**_**i**_ using Coordinate Hit-and-Run with Rounding (CHRR) [3]. However, this is computationally prohibitive for models with millions of thermodynamically realistic orthants. We therefore approximate the distribution in flux space by selecting *n*_*o*_ random orthants according to their relative probability. For each orthant, we then use CHRR to draw a number of flux samples proportional to its probability.

### 2.5 Data and models

We validated our methods on data for *E. coli* growing on different carbon sources [22] with models based on iML1515 [23]. The dataset contains measured growth rates, uptake and secretion rates for 10 metabolites, absolute concentrations for 42 metabolites, and 13C estimates for 26 fluxes in core metabolism. Measurements have been performed in minimal media with eight different carbon sources. Because iML1515 did not allow for solutions satisfying the measured rates on galactose and gluconate, we only performed simulations on acetate, fructose, glucose, glycerol, pyruvate and succinate.

We generated models in three steps: (i) We used *NetworkReducer* ‘s lossy reduction [24] to simplify the subsystems related to lipids and cofactors metabolism. (ii) We applied the results of the thermodynamic curation and we integrated condition-specific constraints such as media composition and measured growth, uptake and secretion rates. The distribution of each intracellular metabolite was set to the log-normal distribution of all metabolites in the acetate and glucose conditions. We used the same mean, but a 5-fold larger standard deviation, for periplasmic and extracellular metabolites (except the ones provided in the media, for which we used the actual concentrations). Measured intracellular concentrations were added only for thermodynamic assessment and when specified in Results. (iii) We applied *NetworkReducer* ‘s lossless compression and manual simplifications to obtain a reduced model. We focused on keeping carbon, amino acid and nucleotide metabolism untouched; hence we will refer to the generated models as iML1515-CAN. The models used for TMFA simulations and uniform sampling were generated with the same approach, but without integrating the results of thermodynamic curation. For PTA and TMFA simulations, the set of thermodynamically constrained reactions Γ consisted of all reactions except for biomass, exchange, water transport and export of all metabolites other than CO2. The average dimensions of the models were *n* = 877, *m* = 501 and *γ* = 717.

## 3 Results

### 3.1 Thermodynamic assessment of iML1515-CAN

First, we searched for inconsistent irreversibilities in our reduced version of the *E. coli* model iML1515 ([23]; see Methods). Seven irreversible reactions had to be set to reversible to make the model thermodynamically feasible. Manual inspection of the standard reaction energies confirmed that all these reactions could be reversed at physiological conditions. For example, we identified an internal cycle between propionate metabolism and the TCA (Fig. 2A) that has no solution unless at least one reaction is reversed or removed.

**Figure 2:**
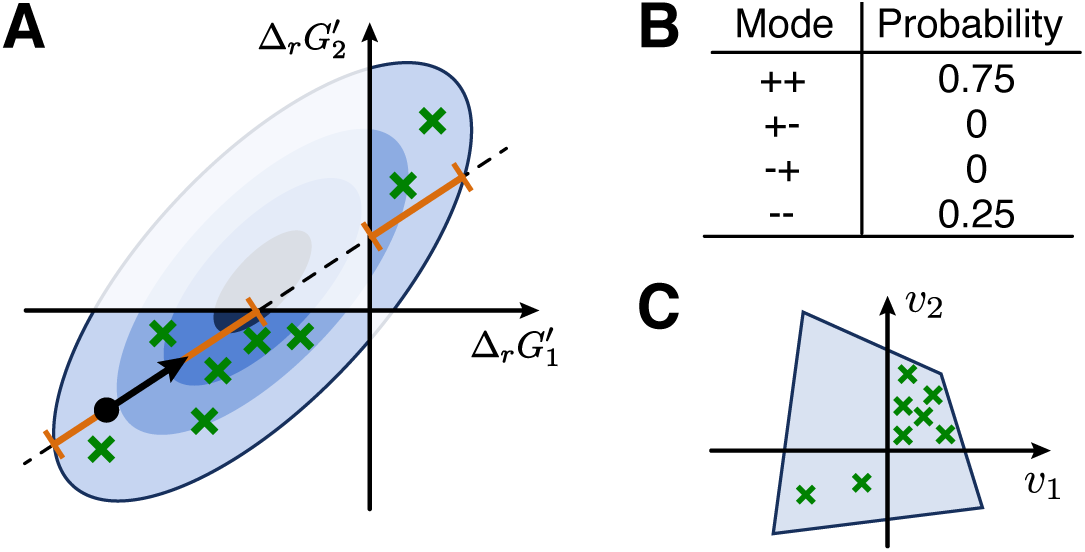
Overview of TFS. (**A**) We sample the thermodynamic space using a modified Hit-and-Run algorithm. At each step, we limit sampling to portions of the space that allow steady state flux distributions (orange). (**B**) As a result, we estimate the probability of each mode. (**C**) Finally, we sample the flux space drawing from each mode a number of samples proportional to its probability.

After resolving the reversibility conflicts, we searched for quantitative anomalies. We selected one glycolytic (glucose) and one gluconeogenic (actetate) growth condition and applied PMO to the corresponding models constrained by the measured concentrations.

We identified 36 unique anomalies in the two models, summarized in Table 1 and detailed in Supplementary Information. The most common reason for anomalies (16 metabolites) is the possible occurrence of *substrate channeling*. Thermodynamics-based modeling assumes a well-mixed system. If two or more enzymes catalyzing consecutive steps of a pathway assemble in a complex, reaction products may find themselves close to the binding pockets of the subsequent enzymes, which increases the reaction probability. In such cases, the intermediate is said to be *channeled* between the two enzymes, breaking the assumption of homogeneity. As a result, reactions that would require unusually high or low concentrations of a reactant can become feasible even at unfavourable intracellular concentrations. An example highlighted by PMO are the first two steps of proline synthesis, *ProB* and *ProA* (Fig. 2D). Constrained by the measured concentrations of glutamate and energy cofactors, PMO predicts a best-case maximum concentration of 14*μ*M for the intermediate glutamate-5-phosphate, 1.4 standard deviations below the mean estimate. Thus, we hypothesized that glutamate-5-phosphate is channeled between *ProB* and *ProA*, relieving the thermodynamic constraint on its intracellular concentration, which is consistent with literature [25]. Consequently, we updated the model by replacing the two steps with a single net reaction.

**Table 1:** Summary of the anomalies in the concentrations predicted by PMO and the resulting modifications to the model in the curation (third column: number of reactions added, removed or converted to lumped reactions). Percentage values are relative to *m* and *γ*, respectively.

| Reasons for anomalies | Anomalies | Reactions |
| --- | --- | --- |
| Substrate channeling with literature support | 9 | 16 |
| Substrate channeling in gluconeogenesis | 3 | 10 |
| Substrate channeling without literature support | 4 | 0 |
| Incomplete stoichiometry | 2 | 2 |
| Inaccurate irreversibility annotations | 6 | 9 |
| Inaccurate group contribution estimates | 4 | 3 |
| Artifacts of steady state constraints | 5 | 5 |
| False positives | 3 | 0 |
| Total | 36 (7.2%) | 45 (6.2%) |

We predicted and found literature support for substrate channeling in several other pathways (see Supplementary Information for details). Most interestingly, with available metabolomics data, four steps of gluconeogenesis (phosphoglycerate mutase, phosphoglycerate kinase, triose-phosphate isomerase and fructose-biphosphate aldolase) appear unfavourable, suggesting enzyme complexes with more favourable steps (phosphoenolpyruvate carboxykinase and fructose-bisphosphatase) that promote substrate channeling. This is consistent with 13C and kinetic studies showing channeling of glycolytic compounds [26, 27].

In addition, two anomalies revealed that the first step of serine synthesis, phosphoglycerate dehydrogenase, is unfavourable with NAD as cofactor. It is known that a more complex mechanism couples the oxidation of 3-phosphoglycerate to the reduction of ubiquinone, which is more favourable [28]. We attributed 15 anomalies to inaccurate Δ_*r*_*G*^°^ estimates, inaccurate irreversibility annotations and cases in which dilution effects likely dominate the steady state constraint. Only 3 anomalies were false positives in which concentrations far from the average are expected, as for intracellular oxygen. Overall, 91.7% of the metabolites flagged after PMO reflected model inaccuracies or knowledge gaps, leading to the curation of 45 reactions, 6.2% of the reactions with thermodynamic constraints.

### 3.2 Sampling modes and directions in iML1515-CAN

We used TFS to sample the thermodynamic space of iML1515-CAN in different growth conditions, both with (*M* +, all conditions) and without (*M*−, all conditions except glucose and acetate, where metabolomics data have been used to curate the model) integrating metabolomics data. Specifically, we collected 10^8^ samples of orthants and 10^5^ samples of **Δ**_**r**_**G**′. In each condition, PMO successfully found initial points and the sampler reached convergence to the target distribution according to common metrics (see Methods). Interestingly, TFS only needed 401 variables to represent **Δ**_**r**_**G**′ of the 717 reactions in Γ. In models constrained only with measured growth and exchange rates, we discovered 16.5·10^6^ orthants, out of which 11.5·10^6^ suffice to cover 95% of the thermodynamic space. These numbers decreased to 11.3 · 10^6^ and 7.2 · 10^6^ once we integrated measured metabolite concentrations. Overall, this represents a substantial reduction over the average theoretical maximum number of orthants (computed from the number of reversible reactions) in the TMFA-constrained models (*M*−: 5.8 · 10^22^, *M* +: 3.1 · 10^20^). Hence, correlations in thermodynamic space may impose important constraints on possible combinations of flux directions.

We compared the predicted reaction directions of TFS, TMFA, and US on TMFA-constrained models and validated them against 13C estimates. For TFS and US, a prediction was considered irreversible when at least 95% of the samples represented the same direction. On average, TMFA could only constrain a small fraction of the models’ 71–79 reversible reactions, in particular without metabolomics data (Fig. 4A). US yielded the most constrained predictions, while TFS predicted irreversibilities for about half of the 102–105 reversible reactions in the PMO-curated models. Despite having the highest precision, US also showed the lowest accuracy: it incorrectly predicted the direction of part of the 11 − 15 reversible fluxes with available 13C estimates. Incorrect predictions focus on two main pathways in gluconeogenic conditions: upper glycolysis, where US systematically prefers fructose 6-phosphate aldolase over fructose-bisphosphate aldolase, and the non-oxidative pentose phosphate pathway. In one condition, these choices result from TMFA incorrectly predicting fructose-bisphosphate aldolase to be irreversible. In all conditions, the irreversibilities predicted by TFS were consistent with the 13C estimates.

**Figure 3:**
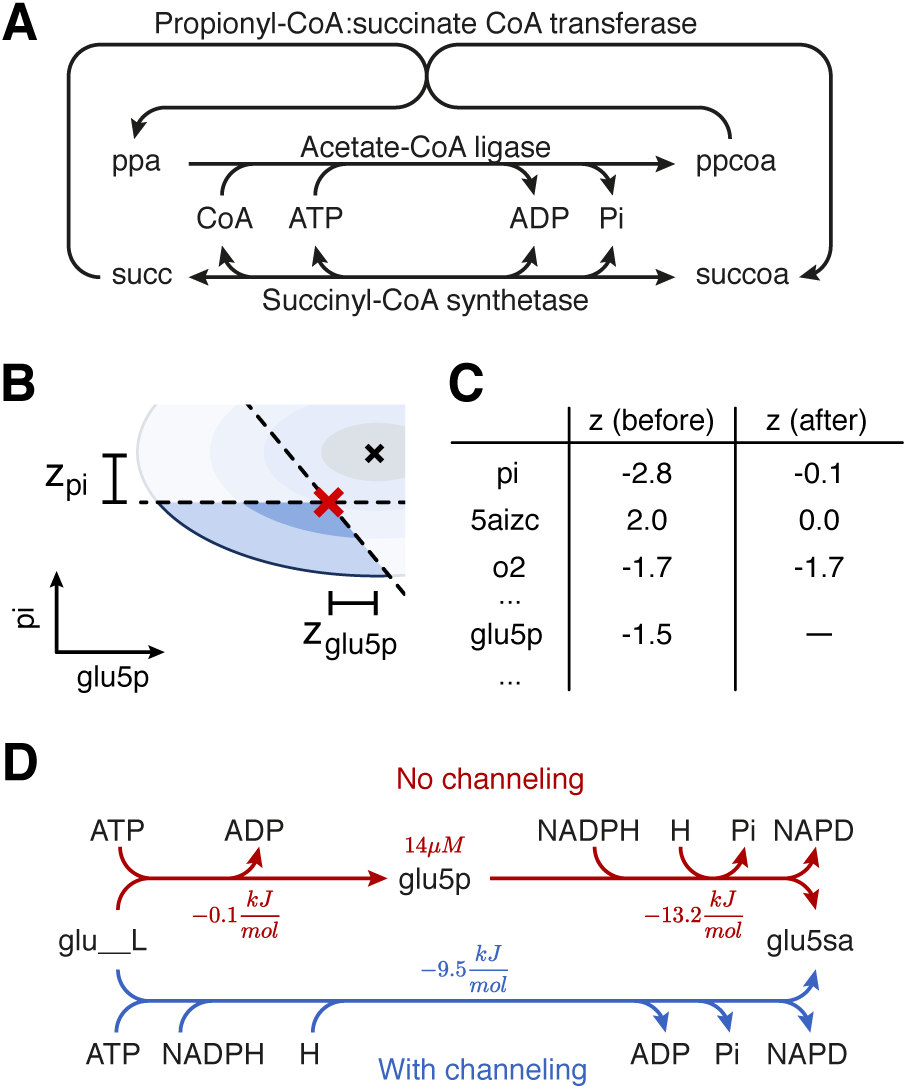
Thermodynamic assessment of iML1515-CAN. (**A**) Example of conflicting irreversibilities. The irreversibility of propionyl-CoA:succinate CoA transferase and acetate-CoA ligase implies that the conversion of succinate to succinyl-CoA is thermodynamically favourable, in conflict with the direction of succinyl-CoA synthetase, which must produce succinate as part of the TCA. (**B**) After using PMO to find **t**^*∗*^ (red cross), we compute the *z*-scores of each metabolite and metabolites, i.e. the normalized distance from the mean (black cross). (**C**) Metabolites with high *z*-score (“before”) are selected. After literature search, 29 out of the 36 flagged by the analysis led to improvements in the model, as shown by smaller *z*-scores in the curated model (“after”). (**D**) A symptomatic case suggesting substrate channeling. An unfavorable reaction produces an intermediate (glutamate-5-phosphate) that must have low concentration to maintain thermodynamic feasibility and is followed by a favorable reaction. The total reaction energies with and without channeling are not identical because the estimated concentrations differ.

**Figure 4:**
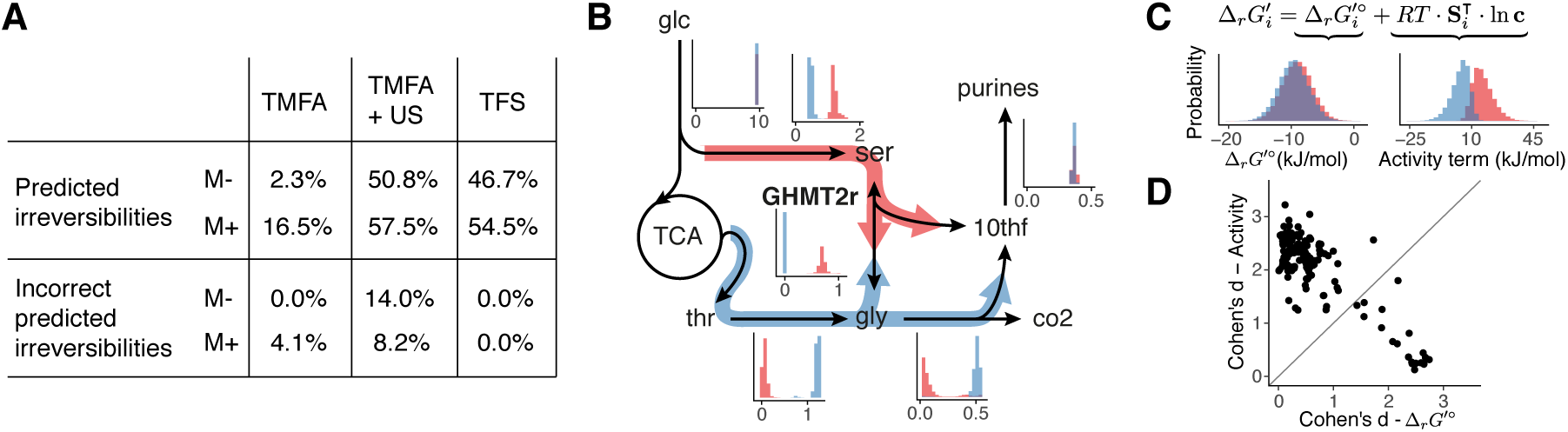
Reaction directions and fluxes. (**A**) Precision and accuracy of predicting directionalities with different methods. Predicted irreversibilities are given as percentage of the average number of reversible reactions in the different conditions. The percentage of incorrectly predicted irreversibilities is relative to the average number of fluxes that are reversible in the models and have 13C estimates. (**B**) Example of multimodal predictions of fluxes with TFS for the synthesis of purines and glycine. Orange: serine is converted to glycine, donating one carbon to form 10-formyltetrahydrofolate (10thf), a cofactor for purine synthesis. Blue: glycine is synthesized from threonine and is converted to CO2 and 10thf. Histograms show predicted flux distributions, clustered by the direction of GHMT2r. (**C**) Distribution of **Δ**_**r**_**G**^′°^ (left) and the activity term (right) of glycine hydroxymethyltransferase (GHMT2r), clustered by the direction of the reaction. 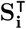 is the *i*-th row of 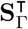. (**D**) Cohen’s d for the distributions of the standard reaction energies and of the activity terms in the two clusters for all reactions in Γ.

Beyond the validated fluxes, we found cases (*M*−: 11 − 18, *M* +: 9 − 13) where both US and TFS predicted a reaction to be irriversible, but in different directions, mostly for reactions that can participate in internal cycles and when alternative paths exist for the synthesis of the same compound. For example, 5-phospho-alpha-D-ribose 1-diphosphate is generally synthesized directly from ribose 5-phosphate (as predicted by TFS), but US predicts a conversion in the opposite direction and 5-phospho-alpha-D-ribose 1-diphosphate synthesis through a longer and stoichiometrically equivalent path via ribose 1-phosphate and ribose 1,5-diphosphate. While literature suggests that the directions predicted by TFS are correct (see Supplementary Information), the available data does not allow for a systematic validation. These differences highlight the effect of using different spaces to determine reaction directions. In US, the probability of a direction is determined by the fraction of flux space that allows it; this favors directions that impose minimal constraints on the rest of the network. In contrast, TFS predicts directions based on the probability of having a favorable reaction energy.

### 3.3 Exploration of metabolic modes

Using the samples of orthants and their weights we sampled the flux space of 10^4^ orthants for each condition. TFS predicted millions of modes in iML1515-CAN, but focusing on the predicted flux distribution of individual reactions one observes fewer than 100 *marginal modes*, which have simpler interpretation. An example are the two modes defined by the direction of glycine hydroxymethyltransferase (GHMT2r) (Fig. 4B). Physiologically, serine is used to synthesize glycine and feed 1C-metabolism, required for purine synthesis. US only predicts this path. TFS predicts an additional mode where glycine and 1C are synthesized from threonine. This *threonine bypass* is not commonly used by *E. coli*, likely because of higher resource requirements. However, it has been successfully used to enhance PHB production [29]. Interestingly, most reactions in glycolysis and the TCA display several marginal modes, depending on the direction of reactions in the rest of the network. While the effect of reversing a reaction on the fluxes can be analyzed with classical FBA or US, TFS provides additional information: the probability (and feasibility) of each direction and the correlation between the direction and the thermodynamic quantities. In the case of GHMT2r, the two modes can be better distinguished by the activity term—more precise estimates of **Δ**_**r**_**G**^′°^ would contribute little information about the reaction direction (Fig. 4C). Conversely, knowing the direction of the reaction would constrain the ratio of serine and glycine, and thus their concentrations. We computed Cohen’s d for the distribution of **Δ**_**r**_**G**^′°^ and of the activity term in the two modes for each reversible reaction (Fig. 4D). In most cases, the modes are better separated by the activity term, suggesting that uncertainties in the standard reaction energy estimates have limited impact. Additional measurements of absolute metabolite concentrations will likely constrain more reaction directions.

### 3.4 Prediction of metabolite concentrations

Finally, we compared the metabolite concentrations predicted by TFS and TMFA to the experimentally measured values in the four *M*− conditions. TFS predictions generally agree with the measured values and highlight strong thermodynamic constraints in the concentrations of some metabolites, such as the NAD/NADH ratio and organic phosphate (Fig. 5A). The mean concentrations predicted by TFS correlate with the experimental measurements (*r* = 0.32, Fig. 5B). For several metabolites, the predicted distribution was very close to the prior distribution (KL smaller than 0.2), suggesting that they participate in reactions that are always favorable. Reducing the coverage of the predictions to metabolites with KL *≥* 0.2 led to a higher correlation (*r* = 0.63), showing that their concentration is thermodynamically constrained. When we focused on the 95% Confidence Interval (CI) of the samples and compared them to the ranges predicted by TMFA (with 95% CI on the distributions of **Δ**_**r**_**G**^′°^ and ln **c**), TFS always predicted narrower ranges (Fig. 5C), with an average 4.5-fold reduction (0.65 on the log_10_-scale). We quantified the overlap of the predicted distributions (uniform within the predicted interval for TMFA) with the distribution of the measurements using the Hellinger distance (Fig. 5D). In 66% of the cases, TFS agreed better with the measurements than TMFA, further supporting its overall higher predictive power for metabolite concentrations.

**Figure 5:**
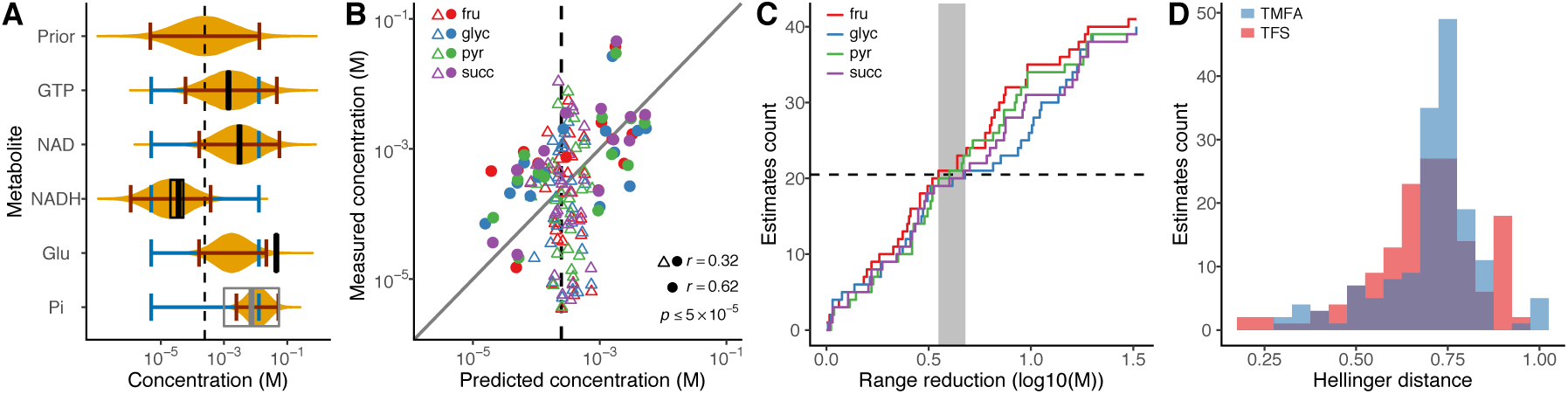
Metabolite concentrations. (**A**) Examples of distributions predicted by TFS (orange), the respective 95% confidence intervals (red), ranges predicted by TMFA (blue) and mean and 95% confidence intervals of the metabolomics measurements (black, range from literature in gray). (**B**) Mean concentrations predicted by TFS agree with measured values (Pearson’s *r* = 0.32, Root Mean Square Error RMSE = 0.87 in log_10_ scale). Reducing the coverage to metabolites with a Kullback-Leibler divergence (KL) between prior and posterior distribution above 0.2 (full circles) increases the agreement to *r* = 0.63, RMSE = 0.72. (**C**) Cumulative distribution of the reduction in the width of the predicted interval of TFS over TMFA for the validated metabolites. (**D**) Hellinger distances between the distribution of the predicted and measured metabolite concentrations for TMFA and TFS.

## 4 Discussion

Realistic models must obey the laws of thermodynamics: quantitative observations inform on the model structure, and mechanistic knowledge constrains parameters. Here, we show how this connection can be exploited in both directions to analyze metabolism. Our PTA framework for thermodynamic-based modeling includes optimization and sampling methods for the evaluation and exploration of metabolic networks. As demonstrated for *E. coli*, it allows to combine metabolic models with experimental data to discover model inaccuracies and generate hypotheses on mechanisms such as substrate channeling. Conversely, we showed that the stoichiometry of a network, coupled with thermodynamic constraints and few physiological observations can significantly reduce the space of feasible metabolic modes and metabolite concentrations.

In contrast to other methods, PTA interprets uncertainties in the thermodynamic parameters probabilistically to account for correlations between uncertainties of multiple estimates. Moreover, we require all reactions to have a well-defined direction. The model cannot silently resolve thermodynamic inconsistencies by setting a flux to zero as in TMFA—problems are visible to the modeler, guiding model curation and the formulation of novel hypotheses. Using the thermodynamic space as prior for sampling the flux space changed the exploration principle from the one used by US. Instead of using the volume of the flux space only, TFS determines the probability of a mode based on the probability of having a set of reaction energies allowing it. This excludes thermodynamically infeasible cycles from the solution. It also prevents distributed thermodynamic bottlenecks [30] and highlights feasible modes that would be missed by US, such as the threonine bypass.

However, PTA comes at the cost of higher computational complexity. Enforcing non-zero flux for each reaction makes optimization problems more difficult to solve, and sampling the non-convex thermodynamic space requires long simulation times; the asymptotic complexity is potentially exponential. While we successfully sampled models with up to 880 reactions, we anticipate that computational advancements are required to work with larger models, for example by modeling different parts of the network independently.

To discover new biology, PMO proved effective at identifying occurrences of substrate channeling. While TMFA suggested such occurrences [17], predictions were limited to cases where predicted concentrations in an essential pathway exceed physiological ranges. Most of our predictions were supported by at least another source in *E. coli* or other organisms, and we also obtained novel hypotheses. That gluconeogenesis appears unfavourable is puzzling: several observations suggest channeling of glycolytic compounds. Yet, this observation could be attributed to inaccurate estimates of standard reaction energies, for example, because magnesium ions [31] and temperature [13], which affect the thermodynamics of glycolysis, are insufficiently accounted for. Additionally, the analysis showed that the curation of reaction directions for FBA applications leads to thermodynamic inaccuracies. As coverage and precision of standard reaction energies and metabolite concentrations increases, we therefore encourage to model thermodynamics explicitly.

Finally, while TFS predicts significantly fewer modes than contained in the flux space, these modes cover a wide range of behaviors. Thermodynamics alone does not determine what a cell does, but it restricts what options are available to achieve a certain phenotype. The final choice probably follows from other constraints or principles such as enzyme cost and minimal resource usage, whose integration could further increase the predictive power of thermodynamically constrained models.

## Acknowledgements

We thank Elad Noor for the help using *equilibrator-api*, Axel Theorell for comments on the manuscript, and Fabian Rudolf for helpful discussions.

