## Supplementary Information for "Probabilistic Thermodynamic Analysis of Metabolic Networks"

### 1 Methods

#### 1.1 Probabilistic Metabolic Optimization (PMO)

We solved PMO problems using the Gurobi solver (v9.0.1) [1]. Eq. (1) summarizes the constraints:

$$\begin{gathered} \mathbf{t} := \begin{bmatrix} \ln \mathbf{c} \\ \mathbf{\Delta_r G'^{\circ}} \\ \mathbf{\Delta_r G'} \end{bmatrix} = \mathbf{Qm} + \mu_{\mathbf{t}} \\ \|\mathbf{m}\| \leq \chi^2_n(\alpha) \\ \mathbf{S} \cdot \mathbf{v} = 0 \\ \mathbf{lb} \leq \mathbf{v} \leq \mathbf{ub} \\ M_f \mathbf{d} - \mathbf{v_{con}} \leq \mathbf{1} \cdot (M_f - \epsilon_f) \\ M_f \mathbf{d} - \mathbf{v_{con}} \geq \mathbf{1} \cdot \epsilon_f \\ M_r \mathbf{d} + \mathbf{\Delta_r G'} \leq \mathbf{1} \cdot (M_r - \epsilon_r) \\ M_r \mathbf{d} + \mathbf{\Delta_r G'} \geq \mathbf{1} \cdot \epsilon_r \end{gathered} \tag{1}$$

Here, the vector of integers $\mathbf{d}$ in $\{0,1\}^{\gamma}$ denotes if a reaction has positive (1) or negative (0) flux and the relationship between free energies and fluxes has been rewritten in the *Big-M* formulation [2]. $M_f$ ($M_r$) are the big-M values and $\epsilon_f$ ($\epsilon_r$) are the minimum magnitudes for fluxes (reaction energies). These values must be chosen carefully as *Big-M* constraints are often a source of numerical errors. In most cases, the range of coefficients should not span more than 9 orders of magnitude. We choose $M_r = 1000\frac{kJ}{mol}$ as it is larger than the $\Delta_r G'^{\circ}$ of any reaction in the model and $\epsilon_r = 0.1\frac{kJ}{mol}$ since smaller values imply high enzyme cost and are thus unlikely. For fluxes, we set $M_f$ larger than the highest flux magnitude (1000 in iML1515-CAN) and $\epsilon_f = 10^{-8} \cdot M_f$. Additionally we set the Gurobi parameters `FeasibiltyTol` $= 10^{-9}$ and `IntFeasTol` $= 10^{-9}$ to ensure that sign constraints are not violated.

The objective depends on the application: for quantitative assessment of the models we minimize $\|\mathbf{m}\|_2$ (i.e. we maximize the probability of $\mathbf{m}$), while for searching initial points for the sampler we maximize and minimize reaction fluxes. We used $\alpha = 0.95$ as confidence level.

The matrix $\mathbf{Q}$ and the vector of $z$-scores $\mathbf{z}$ of a solution $\mathbf{m}$ are computed from the truncated eigenvalue decomposition of $\mathbf{\Sigma_t}$

$$\mathbf{\Sigma_t} = \mathbf{U\Lambda U}^{\intercal} \approx \mathbf{U_q \Lambda_q U_q}^{\intercal} = \mathbf{U_q \Lambda_q}^{1/2} \cdot \left(\mathbf{U_q \Lambda_q}^{1/2}\right)^{\intercal} , \tag{2}$$

where $\mathbf{U}$ is an orthogonal matrix containing the eigenvectors of $\mathbf{\Sigma_t}$ and $\mathbf{\Lambda}$ is a diagonal matrix containing the eigenvalues of $\mathbf{\Sigma_t}$. $\mathbf{U_q}$ and $\mathbf{\Lambda_q}$ are submatrices of $\mathbf{U}$ and $\mathbf{\Lambda}$ that represent eigenvectors and eigenvalues for which the eigenvalue is larger than $10^{-3}$. Given the decomposition, $\mathbf{Q} = \mathbf{U_q \Lambda_q}^{1/2}$ and $\mathbf{z} = \mathbf{U_q m}$.

#### 1.2 Thermodynamics and Flux Sampling (TFS)

The algorithm for sampling $\mathcal{T}$ (see main text) relies on the ability to quickly verify whether an orthant in thermodynamic space is *feasible*, i.e. there is a steady state flux solution that satisfies the directionality constraint of the orthant. We achieve fast feasibility verification with the following optimizations:

1. First, we test a simple condition that is necessary but not sufficient at steady state: for each metabolite, there must be at least one in-flux and one out-flux. This is the case when the system of equations

$$|\mathbf{S} \cdot sign(\mathbf{v})| < \sum_j |\mathbf{S_j}| , \tag{3}$$

where $\mathbf{S_j}$ are the columns of $\mathbf{S}$, is satisfied.

2. We maintain a fixed-size cache ($10^5$ entries) with Least Recently Used (LRU) policy that stores the results of the linear program. If a set of directions has been already tested and the result is present in the cache, we use the result directly without calling the solver.

3. If Eq. (3) and the cache cannot determine the feasibility of an orthant, we verify feasibility with a linear program

$$\begin{aligned} \max_{\mathbf{v}} \quad & \mathbf{0} \\ \text{s.t.} \quad & \mathbf{S} \cdot \mathbf{v} = 0 \\ & \mathbf{lb_{orth}} \leq \mathbf{v} \leq \mathbf{ub_{orth}} \ , \end{aligned} \tag{4}$$

where the $\cdot_{orth}$ subscript indicates the original flux bounds restricted to the directions of the orthant. The optimization problem can be solved quickly because (1) setting the objective to zero, the solver only needs to find one feasible point instead of the optimum and (2) we reuse the same model object for each orthant and only modify the flux bounds. This way the solver can use information from the previous solution.

For each simulation on iML1515-CAN we run 200 chains (starting from different modes of the thermodynamic space) for $2 \cdot 10^8$ steps. The first half of the random walk is treated as warm-up and discarded. From the remaining steps, we collect at regular intervals a total of $10^5$ samples of $\mathbf{\Delta_r G'}$ and $10^8$ direction samples. We then use the samples of $\mathbf{\Delta_r G'}$ to conditionally sample metabolite concentrations and $\mathbf{\Delta_r G'^{\circ}}$. However, due to memory requirements it is more efficient to characterize the probability of each orthant using the signs of the reversible reactions only. These are stored efficiently in a hashmap were the keys are binary serializations of the sign pattern of the reversible reactions and the values indicate the number of times an orthant was sampled.

#### 1.3 Models

The condition-specific models used in the analysis (iML1515-CAN) were generated with the following steps:

1. We used *NetworkReducer*'s lossy reduction [3] allowing removal of the reactions, metabolites in the following subsystems: *Cell Envelope Biosynthesis*, *Glycerophospholipid Metabolism*, *Lipopolysaccharide Biosynthesis / Recycling*, *Membrane Lipid Metabolism*, *Cofactor and Prosthetic Group Biosynthesis*, *Folate Metabolism*, *Murein Biosynthesis*, *Murein Recycling*. Additionally, we allowed for removal of transport and exchange reactions for metabolites that are present exclusively in the subsystems above. During reduction, we protected the model's capability of reproducing the measured growth and exchange rates. Reducing the level of detail in these subsystems was necessary to make the model computationally tractable and to ignore parts of the network where the thermodynamic model could be unreliable (e.g. different phase and poorly defined metabolites in lipid metabolism).

2. We manually removed/lumped reactions that only participate to secretion of metabolites or energy-wasting loops in the subsystems above.

3. We removed reactions related to oligosaccharides (glycogen and maltose) as their production and degradation form unfeasible cycles at steady state and the annotated directionalities are thermodynamically inaccurate.

4. PFK_3 and FBA3 were removed to make fluxes in the pentose phosphate pathway comparable to the 13C estimates.

5. We integrated experimental data (metabolite concentrations when required and measured growth/exchange rates).

6. The model was further reduced using *NetworkReducer* lossless compression. We protected all reactions in carbon, amino acid and nucleotide metabolism, which we believed were the most important in the growth conditions (minimal media).

7. As we require non-zero flux through all reactions modeled with thermodynamic constraints, we removed all blocked reactions.

The resulting models maintain an intact description of carbon, amino acid and nucleotide metabolism. Intracellular metabolomics data were only used for model assessment (glucose and acetate conditions) and in the $M+$ conditions. In all conditions we set extracellular concentrations according to the composition of M9 media.

#### 2 Thermodynamic assessment of iML1515-CAN

Table SI 1 lists the irreversible reactions that we had to make reversible to avoid thermodynamic inconsistencies. The same inconsistencies were found in all growth conditions. Table SI 2 shows the anomalies found for the growth on glucose and acetate, together with the explanation we found based on literature and the curation steps. We curated the model only in presence of support from literature. It is possible that some of the irreversibility annotations that we removed during the curation process were added to the model because it is known that *E. coli* does not actively use these reactions in the opposite direction. This is an intentional choice, as thermodynamic reversibility and regulatory choices are two different kinds of constraints. While knowledge of the preferred direction of a reaction can be added on top of a thermodynamically constrained model (e.g. for further reducing the number of modes predicted by TFS), this should not be used as a thermodynamic constraint.

Solving the PMO optimization problem took a time varying between 1 and 4 minutes on a Intel® i7-8700K processor depending on the growth condition and whether the results of the curation were applied or not.

Table SI 1: List of the irreversible reactions in iML1515-CAN that we had to make reversible to avoid thermodynamic inconsistencies.

| Reaction | Reason |
|---|---|
| ACCOAL | Enforces unfeasible internal cycle. |
| ACt4pp | Conflicts with ACt2rpp and NAt3pp. |
| GLYCLTt4pp | Conflicts with GLYCLTt2rpp and NAt3pp. |
| PPAt4pp | Prevents excretion of propionate, enforcing an unfeasible internal cycle. |
| PPCSCT | Enforces unfeasible internal cycle. |
| PROt4pp | Conflicts with PROt2rpp and NAt3pp. |
| PTA2 | Enforces unfeasible internal cycle. |

Table SI 2: Interpretation of the PMO results. NA denotes values that are not available because the metabolite was removed after model curation.

| Metabolite | *z*-score | | Concentration (mM) | | Explanation |
|---|---|---|---|---|---|
| | Before | After | Before | After | |
| Metabolites with $\lvert z \rvert \geq 1$ for growth on glucose. | | | | | |
| pi_c<br>pi_e | −2.8<br>1.4 | −0.1<br>0.0 | 14.2<br>64.0 | 24.6<br>55.7 | Import of the default phosphate specie ($HPO_4^{2-}$) through PIt2rpp is thermodynamically unfeasible at the given concentrations. Phosphate must be transported either as $H_2PO_4^-$ or through PIuabcpp. **Action:** Make PIt2rpp reversible and add reaction for the transport of $H_2PO_4^-$ [4]. |
| mqn8_c<br>mql8_c | 2.4<br>−2.4 | 0.8<br>−0.8 | 30.4<br>0.002 | 1.3<br>0.05 | The reactions of the respiratory complex I have been found to be reversible in mitochondria [5]. It is likely that reversibilities apply also in the bacterial orthologs. **Action:** Make the reactions NADH16pp, NADH17pp and NADH18pp reversible, as their irreversibility annotation is thermodynamically inaccurate. |
| aspsa_c | −2.4 | NA | 0.002 | NA | Substrate channeling has been shown experimentally between ASPK, ASAD and HSDy [6]. We hypothesize channeling through ASPK, ASAD and DHDPS as the respective enzymes are predicted to bind [7]. **Action:** Convert ASPK, ASAD, HSDy and DHDPS to the lumped reactions ASPK_ASAD_HSDy and ASPK_ASAD_DHDPS. |
| 5caiz_c<br>5aizc_c | 2.0<br>−2.0 | NA<br>0.0 | 14.6<br>0.004 | NA<br>0.25 | We hypothesize channeling of 5caiz_c between AIRC2 and AIRC3, since the respective enzymes purK and purE have been shown to bind in *E. coli* [7]. Note that this is controversial [8]. **Action:** Convert AIRC2 and AIRC3 to the lumped reaction AIRC2_AIRC3. |
| o2_c | −1.7 | −1.7 | 0.008 | 0.008 | Expected. Limited by extracellular o2 concentration. **Action:** None. |
| 3pg_c<br>3php_c | 1.6<br>−1.6 | 0.26<br>−0.26 | 6.4<br>0.010 | 0.42<br>0.15 | PGCD alone is thermodynamically unfavorable. SerA has been shown to couple 3pg dehydrogenation to akg reduction in *Pseudomonas*. The same mechanism is likely employed by *E. coli* as well [9]. **Action:** Make PGCD reversible. Add reactions for the proposed mechanism: 3pg_c + akg_c $\rightleftharpoons$ 3php_c + r2hglut_c and akg_c + q8h2_c $\rightleftharpoons$ r2hglut_c + q8_c. |
| acg5p_c<br>acglu_c | −1.5<br>1.5 | 0.0<br>NA | 0.012<br>5.2 | 0.25<br>NA | Channeling of acglu between ACGS and ACGK has been shown experimentally in *S. cerevisiae* [10], we hypothesize a similar mechanism in *E. coli.* **Action:** Convert ACGS and ACGK to the lumped reactions ACGS_ACGK. |
| glu5p_c | −1.5 | NA | 0.012 | NA | Channeling of glu5p from GLU5K to G5SD is experimentally supported in *E. coli* [11]. **Action:** Convert GLU5K and G5SD to the lumped reaction GLU5K_G5SD. |

Table SI 2: Interpretation of the PMO results. NA denotes values that are not available because the metabolite was removed after model curation.

| Metabolite | $z$-score | | Concentration (mM) | | Explanation |
|---|---|---|---|---|---|
| | Before | After | Before | After | |
| 4abut_c<br>sucsal_c | 1.5<br>−1.5 | 0.0<br>NA | 5.1<br>0.013 | 0.25<br>NA | Physical interaction between the enzymes catalyzing ABTA and SSALx has been shown experimentally in mitochondria [12]. We hypothesize a similar interaction in *E. coli* leading to channeling of sucsal for both SSALx and SSALy. **Action:** Convert ABTA, SSALx and SSALy to the lumped reactions ABTA_SSALx and ABTA_SSALy |
| ser__L_c<br>2amsa_c | 1.5<br>−1.5 | 0.0<br>NA | 4.7<br>0.014 | 0.25<br>NA | Artifact of the steady-state constraint. 2amsa is a required intermediate for the synthesis of D-serine from L-serine. However, it is unlikely that *E. coli* actively synthesizes D-serine as it is not involved in any cellular function. **Action:** Remove 2amsa_c and ser__D_c to avoid imposing unrealistic constraints on the concentration of L-serine. |
| succoa_c<br>coa_c<br>sl2a6o_c<br>thdp_c | 1.3<br>−1.2<br>−1.1<br>1.1 | 0.60<br>−0.33<br>0.0<br>NA | 3.2<br>0.0245<br>0.027<br>2.4 | 0.83<br>0.13<br>0.25<br>NA | Inaccuracies in group contribution estimates. DHDPS is a ring-forming reaction synthesizing 23dhdp, which is then converted to thdp. THDPS then opens the thdp ring, forming sl2a6o. Both 23dhdp and thdp are not involved in any reaction in TECRDB, thus their formation energy must be estimated from group contribution, which is known to be unreliable for ring alterations [13]. **Action:** Ignore the ring thermodynamics by replacing DHDPS, DHDPRy and THDPS with the lumped reaction DHDPS_DHDPRy_THDPS. |
| ptrc_c<br>4abutn_c | 1.3<br>−1.2 | 1.1<br>−1.1 | 3.1<br>0.021 | 2.3<br>0.027 | STRING predicts possible binding between PatA an PatD. However, we could not find further evidence supporting substrate channeling between the two enzymes. **Action:** None. |
| gcald_c<br>4hthr_c<br>gly_c | 1.1<br>−1.1<br>1.1 | 0.20<br>NA<br>0.0 | 2.5<br>0.026<br>2.5 | 0.39<br>NA<br>0.25 | As 4hthr does not participate in any essential pathway and its maximum flux is very low, the validity of the constraints from the directions of 4HTHRA and 4HTHRK is questionable. *in-vivo* those may be affected by transport and dilution. **Action:** Remove 4HTHRA and 4HTHRK to avoid imposing unrealistic constraints on the concentration of glycine. |
| acon_c | −1.0 | −1.0 | 0.030 | 0.030 | (1) Citrate dehydratase is an unfavourable reaction, while the previous step (citrate synthase) is predicted to be highly favourable ($\Delta_r G' - 40.5 \frac{kJ}{mol}$), suggesting channeling of citrate between the two reactions. Indeed there is experimental evidence for a mitochondrial enzyme complex including malate dehydrogenase, citrate synthase and aconitase [14, 15]. We hypothesize a similar interaction in *E. coli*, leading to the channeling of oxaloacetate and citrate. (2) It is not clear whether aconitate is channeled between the two aconitase steps, since it must be released from the enzyme and turn 180° before binding again. [16] We conservatively assume it is. **Action:** Add lumped reactions for the potential channels (1) MDH_to_CS, CS_to_ACONT, MDH_to_ACONT and (2) ACONTa_ACONTb . |
| Additional metabolites with concentration $\geq 10mM$ for growth on glucose. | | | | | |
| cl_p | 0.57 | 0.3 | 75.5 | 6.0 | The annotated irreversibility of CLt3_2pp is thermodynamically inaccurate and implies high periplasmic concentrations of cl_L. **Action:** Make CLt3_2pp reversible. |
| glu__L_p | 0.46 | 0.0 | 26.2 | 0.25 | The annotated irreversibilities of GLUt4pp, GLUABUTt7pp and ABUTt2pp are thermodynamically inaccurate and imply high periplasmic concentrations of glu__L. **Action:** Make GLUt4pp, GLUABUTt7pp and ABUTt2pp reversible. |
| Additional metabolites with $\|z\| \geq 1$ for growth on acetate. | | | | | |
| glyc_c<br>g3p_c<br>dha_c | 2.2<br>−1.5<br>1.4 | 0.36<br>−1.7<br>−0.36 | 19.9<br>0.012<br>4.2 | 0.52<br>0.007<br>0.12 | Gluconeogenesis appears unfavourable at the given intracellular concentrations. *in-vivo*, unfavourable reactions could be overcome by substrate channeling (see main text). To make unfavourable reactions feasible, they would need to be coupled with favourable steps (PPCK and FBP). **Action:** Based on evidence from literature and predicted requirements we hypothesised two complexes performing substrate channeling: (1) PPCK, ENO, PGM, PGK, GAPD, TPI (or any subset starting from PPCK) and (2) FBP, FBA, TPI (or any subset starting from FBP). The thermodynamics of glycolysis is highly influenced by magnesium ions [17], currently not accounted for by eQuilibrator. While it is likely that some form of channeling involving FBP and PPCK occurs in the pathway, the exact composition of the enzyme complexes is hard to determine. The predicted complexes are thus possibly larger than the actual ones. |
| ac_c | 1.5 | 1.3 | 4.8 | 3.6 | Plausible given the growth condition. **Action:** None. |
| g1p_c | −1.2 | −1.2 | 0.022 | 0.023 | STRING predicts possible binding between Pgm an Pgi, but we could not find further literature evidence supporting substrate channeling between the two enzymes. Moreover, glucose-6-phosphate isomerase operates close to equilibrium, thus substrate channeling between the two enzymes would not bring any thermodynamic advantage. **Action:** None. |
| sucgsa_c<br>sucorn_c | −1.0<br>1.0 | −0.87<br>0.87 | 0.032<br>2.0 | 0.044<br>1.4 | STRING predicts possible binding between AstC an AstD, however we could not find further literature evidence supporting substrate channeling between the two enzymes. **Action:** None. |

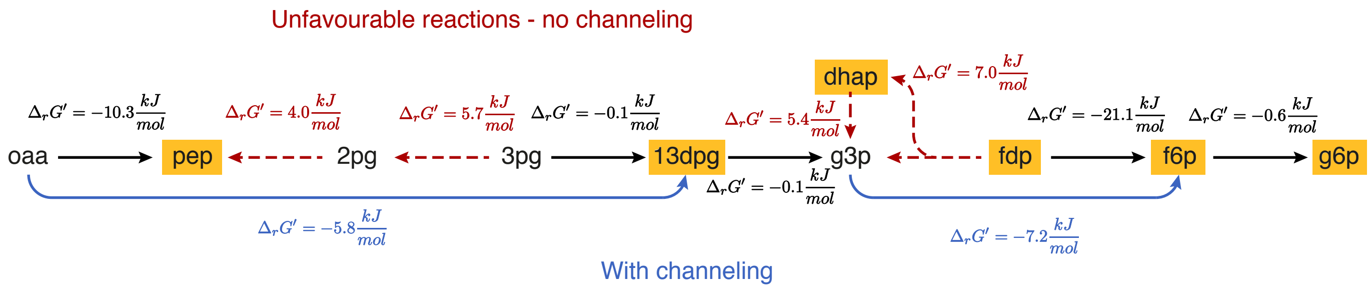


Figure SI 1: Overview of the predicted reaction energies in gluconeogenesis. The concentrations of the metabolites highlighted in orange, as well as all cofactors other than CO2 and phosphate were constrained by measurements. Phosphate is constrained to literature values. Reactions predicted to be unfavorable are shown in red. An example of hypothetical net reactions caused by substrate channeling is shown in blue.

### 3 Sampling iML1515-CAN

#### 3.1 Convergence and performance

For all six conditions, the random walks satisfied recommended convergence metrics [18] (split-R $\leq 1.1$ and $ESS < 5 \cdot n_c$, where $n_c$ is the number of chains). Note that we computed these statistics on the split-chains, meaning that, after discarding the warm-up steps, we split each chain in two halves and treated each half as a chain. This is recommended to detect systematic trends in the chains. After sampling the thermodynamic space, we randomly selected $10^4$ modes according to their probabilities and used a custom C++ implementation of CHRR to sample their flux spaces. The number of flux samples drawn for each orthant was proportional to its probability.

Sampling the thermodynamic space took 10-11 hours depending on the growth condition using 200 Intel® Xeon® CPU E5-2697 cores (year 2013) on our HPC cluster. Uniform sampling of the orthants required additional 5 hours. As a comparison, we also sampled a core model of *E. coli* (109 reactions). There, sampling the thermodynamic and flux spaces took less than 5 minutes on a Intel® i7-8700K processor. Thus, we expect the application of TFS to models containing a few hundred reactions to be computationally affordable without an advanced computational infrastructure. Moreover, the runtime depends mostly on the dimensionality of the thermodynamics space. If an application requires investigation of specific areas of the network only, one can apply thermodynamic constraints only to the corresponding reactions, resulting in faster simulations.

#### 3.2 Prediction of directions and flux distributions

Table SI 3 shows the number of orthants in each condition. These cover a wide range of behaviors, which reflect on the flux distributions of individual reactions (Figure SI 2). Interestingly, Uniform Sampling (US) overlooks many of the capabilities of the network. This is likely an artifact of representing the flux space with a polytope. In high dimensions, most of the volume lies in the center of the space and orthants that only appear close to the boundary of the polytope are too unlikely to be found, independent of their thermodynamic probability.

Table SI 4 summarizes the results of the validation of precision and accuracy of US and TFS against 13C estimates. However, 13C estimates have limited coverage and there were several cases where US and TFS predicted the irreversibility of a reaction but in different directions. We manually validated the predicted directions using EcoCyc [19] (Table 5 and 6) and found that in all verifiable cases TFS predicts the correct direction.

#### 3.3 Prediction of metabolite concentrations

For comparisons against TMFA we used the matTFA [20] implementation, modified to use $\mathbf{\Delta_r G'^{\circ}}$ estimates from eQuilibrator. We constrained the estimates of metabolite concentrations and $\mathbf{\Delta_r G'^{\circ}}$ to their 95% confidence interval. Figure SI 3 shows the predicted TFS distributions and Thermodynamics-based Metabolic Flux Analysis (TMFA) ranges for each metabolite with measured concentration. TMFA could not constrain the concentration of any of those metabolites. This is consistent with the results obtained by the authors of TMFA, which showed constrained metabolite ranges only assuming no error in the standard reaction energies [21].

Table SI 3: Number of orthants found in each condition. The last column shows the minimum number of orthants required to cover 95% of the thermodynamic space.

| Metabolomics | Condition | N. orthants | Mean samples per orthant | % orthants for 95% coverage |
|---|---|---|---|---|
| M- | fru | 17761983 | 5.63 | 71.85% |
| M- | glyc | 13952353 | 7.17 | 66.39% |
| M- | pyr | 17058219 | 5.86 | 70.69% |
| M- | succ | 17099759 | 5.85 | 70.76% |
| M+ | fru | 10092355 | 9.91 | 62.25% |
| M+ | glyc | 8760400 | 11.42 | 60.88% |
| M+ | pyr | 14382468 | 6.95 | 67.01% |
| M+ | succ | 12381754 | 8.08 | 65.49% |
| M+ | ac | 10539012 | 9.49 | 63.12% |
| M+ | glc | 11596903 | 8.62 | 64.58% |
| M- | Average: | 16468079 | 6.13 | 70.11% |
| M+ | | 11292149 | 9.08 | 64.21% |

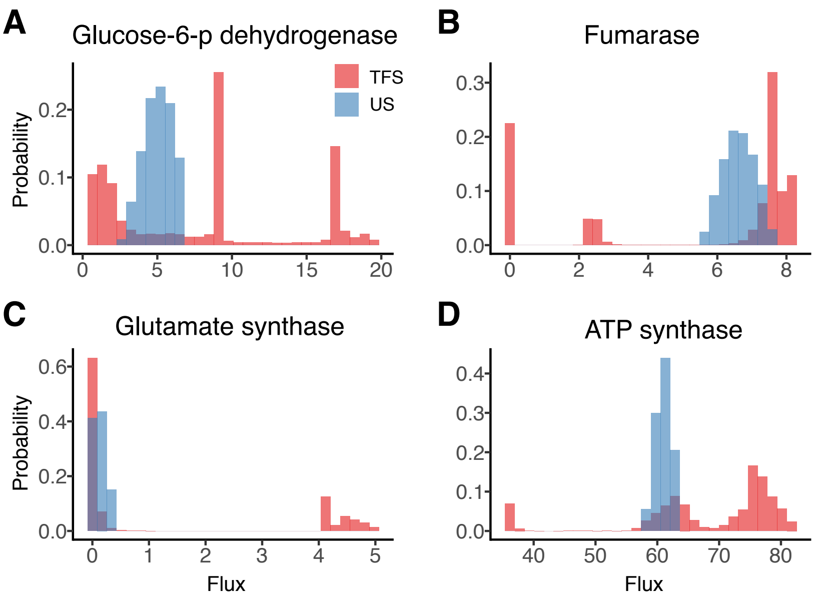


Figure SI 2: Selected examples comparing the flux distributions predicted by US and TFS ($M-$, growth on fructose). (**A**) Large variability in the flux through the pentose phosphate pathway. (**B**) A "branched TCA" (inactive or reverse fumarase) is themodynamically realistic. (**C**) *E. coli* has two paths for synthesizing glutamate. The path through glutamate synthase requires more ATP than the path through glutamate dehydrogenase, but may be the only thermodynamically feasible option in low-nitrogen conditions. (**D**) The flux through oxidative phosphorylation can vary significantly depending on the directions selected in the rest of the network.

Table SI 4: Comparison of precision and accuracy of different methods (see main text). Counts (percentages) are given as range (mean) over all conditions. The last two rows report the number (percentage relative to the reversible reactions) of reactions for which US and TFS predicted irreversibility in different directions.

| | TMFA | | TMFA+US | | TFS | |
|---|---|---|---|---|---|---|
| | M- | M+ | M- | M+ | M- | M+ |
| Rxns in model | 864-870 | 864-871 | 864-870 | 864-871 | 875-881 | 875-883 |
| Reversible rxns in model | 73-79 | 71-79 | 73-79 | 71-79 | 102-105 | 102-105 |
| Reversible fluxes in model | 11-14 | 10-14 | 11-14 | 10-14 | 13-15 | 13-15 |
| Predicted reversible reactions | 71-77 | 57-70 | 32-40 | 28-35 | 53-57 | 46-48 |
| (% of reversible rxns in model) | 97.7% | 83.5% | 49.2% | 42.5% | 53.3% | 45.5% |
| Predicted irreversible reactions (% of reversible rxns in model) | 2.3% | 16.5% | 50.8% | 57.5% | 46.7% | 54.5% |
| Incorrect predicted irreversibilities | 0 | 0-3 | 0-5 | 0-3 | 0 | 0 |
| (% of reactions covered by 13C estimates) | 0.0% | 4.1% | 14.0% | 8.2% | 0.0% | 0.0% |
| Conflicting (TMFA + US vs TFS) | | | | | 11-18 | 9-13 |
| (% of reversible rxns in US model) | | | | | 17.2% | 14.7% |

Table SI 5: Evaluation of the predicted directions of reactions for which US and TFS predicted irreversibilities in opposing directions ($M-$). Predictions that are correct according to EcoCyc [19] are highlighted in green. Light green indicates predictions that we believe are correct but could not be confirmed. We did not seek validation for transporters (gray) as their exact stoichiometry is often unclear and. Moreover, we ignored reactions already validated against 13C data and reactions that are potentially involved in substrate channeling (in that case, the direction of the reaction does not reflect the direction of the net flux).

| Reaction | Condition | US | TFS | Explanation | Reference |
|---|---|---|---|---|---|
| ACCOAL | fru | FW | BW | Not found in ecocyc, DG° favors BW | |
| ACCOAL | glyc | FW | BW | Not found in ecocyc, DG° favors BW | |
| ACCOAL | pyr | FW | BW | Not found in ecocyc, DG° favors BW | |
| ACCOAL | succ | FW | BW | Not found in ecocyc, DG° favors BW | |
| ACt4pp | fru | FW | BW | | |
| ACt4pp | glyc | FW | BW | | |
| ACt4pp | pyr | FW | BW | | |
| ACt4pp | succ | FW | BW | | |
| CBMKr | fru | FW | BW | BW | https://ecocyc.org/ECOLI/NEW-IMAGE?type=PATHWAY&object=PWY0-41 |
| CBMKr | glyc | FW | BW | BW | https://ecocyc.org/ECOLI/NEW-IMAGE?type=PATHWAY&object=PWY0-41 |
| CBMKr | pyr | FW | BW | BW | https://ecocyc.org/ECOLI/NEW-IMAGE?type=PATHWAY&object=PWY0-41 |
| CBMKr | succ | FW | BW | BW | https://ecocyc.org/ECOLI/NEW-IMAGE?type=PATHWAY&object=PWY0-41 |
| CRNDt2rpp | glyc | FW | BW | | |
| G3PD2 | glyc | FW | BW | BW | https://ecocyc.org/ECOLI/NEW-IMAGE?type=REACTION&object=GLYC3PDEHYDROGBIOSYN-RXN |
| GLYAT | fru | FW | BW | BW | https://ecocyc.org/ECOLI/NEW-IMAGE?type=PATHWAY&object=THREONINE-DEG2-PWY |
| GLYAT | glyc | FW | BW | BW | https://ecocyc.org/ECOLI/NEW-IMAGE?type=PATHWAY&object=THREONINE-DEG2-PWY |
| GLYCLTt4pp | fru | FW | BW | | |
| GLYCLTt4pp | glyc | FW | BW | | |
| GLYCLTt4pp | pyr | FW | BW | | |
| GLYCLTt4pp | succ | FW | BW | | |
| NADH17pp | fru | FW | BW | Possibly BW (see thermodynamic assessment) | |
| NADH17pp | glyc | FW | BW | Possibly BW (see thermodynamic assessment) | |
| NADH17pp | pyr | FW | BW | Possibly BW (see thermodynamic assessment) | |
| NADH17pp | succ | FW | BW | Possibly BW (see thermodynamic assessment) | |
| PPM | fru | BW | FW | FW | https://ecocyc.org/ECOLI/NEW-IMAGE?type=PATHWAY&object=PWY0-1295 |
| PPM | glyc | BW | FW | FW | https://ecocyc.org/ECOLI/NEW-IMAGE?type=PATHWAY&object=PWY0-1295 |
| PPM | pyr | BW | FW | FW | https://ecocyc.org/ECOLI/NEW-IMAGE?type=PATHWAY&object=PWY0-1295 |
| PPM | succ | BW | FW | FW | https://ecocyc.org/ECOLI/NEW-IMAGE?type=PATHWAY&object=PWY0-1295 |
| PROt4pp | fru | FW | BW | | |
| PROt4pp | glyc | FW | BW | | |
| PROt4pp | pyr | FW | BW | | |
| PROt4pp | succ | FW | BW | | |
| PRPPS | fru | BW | FW | FW | https://ecocyc.org/ECOLI/NEW-IMAGE?type=REACTION&object=PRPPSYN-RXN |
| PRPPS | glyc | BW | FW | FW | https://ecocyc.org/ECOLI/NEW-IMAGE?type=REACTION&object=PRPPSYN-RXN |
| PRPPS | pyr | BW | FW | FW | https://ecocyc.org/ECOLI/NEW-IMAGE?type=REACTION&object=PRPPSYN-RXN |
| PRPPS | succ | BW | FW | FW | https://ecocyc.org/ECOLI/NEW-IMAGE?type=REACTION&object=PRPPSYN-RXN |
| TRSARr | fru | BW | FW | FW | https://ecocyc.org/ECOLI/NEW-IMAGE?type=REACTION&object=TSA-REDUCT-RXN |
| TRSARr | glyc | BW | FW | FW | https://ecocyc.org/ECOLI/NEW-IMAGE?type=REACTION&object=TSA-REDUCT-RXN |
| TRSARr | pyr | BW | FW | FW | https://ecocyc.org/ECOLI/NEW-IMAGE?type=REACTION&object=TSA-REDUCT-RXN |
| TRSARr | succ | BW | FW | FW | https://ecocyc.org/ECOLI/NEW-IMAGE?type=REACTION&object=TSA-REDUCT-RXN |

Table SI 6: Evaluation of the predicted directions of reactions for which US and TFS predicted irreversibilities in opposing directions ($M+$). Predictions that are correct according to EcoCyc [19] are highlighted in green. Light green indicates predictions that we believe are correct but could not be confirmed. We did not seek validation for transporters (gray) as their exact stoichiometry is often unclear and. Moreover, we ignored reactions already validated against 13C and reactions that are potentially involved in substrate channeling (in that case the direction of the reaction does not reflect the direction of the net flux).

| Reaction | Condition | US | TFS | Explanation | Reference |
|---|---|---|---|---|---|
| ACCOAL | ac | FW | BW | Not found in ecocyc, DG° favors BW | |
| ACCOAL | fru | FW | BW | Not found in ecocyc, DG° favors BW | |
| ACCOAL | glc | FW | BW | Not found in ecocyc, DG° favors BW | |
| ACCOAL | glyc | FW | BW | Not found in ecocyc, DG° favors BW | |
| ACCOAL | pyr | FW | BW | Not found in ecocyc, DG° favors BW | |
| ACCOAL | succ | FW | BW | Not found in ecocyc, DG° favors BW | |
| ACKr | glyc | FW | BW | BW if not growing on acetate. | https://ecocyc.org/ECOLI/NEW-IMAGE?type=PATHWAY&object=PWY0-1312 |
| ACt4pp | fru | FW | BW | | |
| ACt4pp | glc | FW | BW | | |
| ACt4pp | glyc | FW | BW | | |
| ACt4pp | pyr | FW | BW | | |
| ACt4pp | succ | FW | BW | | |
| CBMKr | ac | FW | BW | BW | https://ecocyc.org/ECOLI/NEW-IMAGE?type=PATHWAY&object=PWY0-41 |
| CBMKr | fru | FW | BW | BW | https://ecocyc.org/ECOLI/NEW-IMAGE?type=PATHWAY&object=PWY0-41 |
| CBMKr | glc | FW | BW | BW | https://ecocyc.org/ECOLI/NEW-IMAGE?type=PATHWAY&object=PWY0-41 |
| CBMKr | glyc | FW | BW | BW | https://ecocyc.org/ECOLI/NEW-IMAGE?type=PATHWAY&object=PWY0-41 |
| CBMKr | pyr | FW | BW | BW | https://ecocyc.org/ECOLI/NEW-IMAGE?type=PATHWAY&object=PWY0-41 |
| CBMKr | succ | FW | BW | BW | https://ecocyc.org/ECOLI/NEW-IMAGE?type=PATHWAY&object=PWY0-41 |
| CRNDt2rpp | ac | FW | BW | | |
| CRNDt2rpp | succ | FW | BW | | |
| F6PA | ac | BW | FW | Not specified in ecocyc pathways | |
| F6PA | glyc | BW | FW | Not specified in ecocyc pathways | |
| F6PA | pyr | BW | FW | Not specified in ecocyc pathways | |
| F6PA | succ | BW | FW | Not specified in ecocyc pathways | |
| GLYAT | fru | FW | BW | BW | https://ecocyc.org/ECOLI/NEW-IMAGE?type=PATHWAY&object=THREONINE-DEG2-PWY |
| GLYAT | glc | FW | BW | BW | https://ecocyc.org/ECOLI/NEW-IMAGE?type=PATHWAY&object=THREONINE-DEG2-PWY |
| GLYAT | glyc | FW | BW | BW | https://ecocyc.org/ECOLI/NEW-IMAGE?type=PATHWAY&object=THREONINE-DEG2-PWY |
| GLYCLTt4pp | ac | FW | BW | | |
| GLYCLTt4pp | fru | FW | BW | | |
| GLYCLTt4pp | glc | FW | BW | | |
| GLYCLTt4pp | glyc | FW | BW | | |
| GLYCLTt4pp | pyr | FW | BW | | |
| GLYCLTt4pp | succ | FW | BW | | |
| NADH17pp | ac | FW | BW | Possibly BW (see thermodynamic assessment) | |
| NADH17pp | fru | FW | BW | Possibly BW (see thermodynamic assessment) | |
| NADH17pp | pyr | FW | BW | Possibly BW (see thermodynamic assessment) | |
| NADH18pp | ac | FW | BW | Possibly BW (see thermodynamic assessment) | |
| NADH18pp | fru | FW | BW | Possibly BW (see thermodynamic assessment) | |
| NDPK1 | ac | BW | FW | FW | https://ecocyc.org/ECOLI/NEW-IMAGE?type=REACTION&object=GDPKIN-RXN |
| NDPK1 | fru | BW | FW | FW | https://ecocyc.org/ECOLI/NEW-IMAGE?type=REACTION&object=GDPKIN-RXN |
| NDPK1 | glyc | BW | FW | FW | https://ecocyc.org/ECOLI/NEW-IMAGE?type=REACTION&object=GDPKIN-RXN |
| POR5 | ac | BW | FW | FW, distributed bottleneck with RNTR1*2 | |
| PPM | fru | BW | FW | FW | https://ecocyc.org/ECOLI/NEW-IMAGE?type=PATHWAY&object=PWY0-1295 |
| PPM | glc | BW | FW | FW | https://ecocyc.org/ECOLI/NEW-IMAGE?type=PATHWAY&object=PWY0-1295 |
| PPM | succ | BW | FW | FW | https://ecocyc.org/ECOLI/NEW-IMAGE?type=PATHWAY&object=PWY0-1295 |
| PROt4pp | ac | FW | BW | | |
| PROt4pp | fru | FW | BW | | |
| PROt4pp | glc | FW | BW | | |
| PROt4pp | glyc | FW | BW | | |
| PROt4pp | pyr | FW | BW | | |
| PROt4pp | succ | FW | BW | | |
| PRPPS | fru | BW | FW | FW | https://ecocyc.org/ECOLI/NEW-IMAGE?type=REACTION&object=PRPPSYN-RXN |
| PTAr | glyc | BW | FW | FW if not growing on acetate. | https://ecocyc.org/ECOLI/NEW-IMAGE?type=PATHWAY&object=PWY0-1312 |
| TRSARr | ac | BW | FW | FW | https://ecocyc.org/ECOLI/NEW-IMAGE?type=REACTION&object=TSA-REDUCT-RXN |
| TRSARr | fru | BW | FW | FW | https://ecocyc.org/ECOLI/NEW-IMAGE?type=REACTION&object=TSA-REDUCT-RXN |
| TRSARr | glc | BW | FW | FW | https://ecocyc.org/ECOLI/NEW-IMAGE?type=REACTION&object=TSA-REDUCT-RXN |
| TRSARr | glyc | BW | FW | FW | https://ecocyc.org/ECOLI/NEW-IMAGE?type=REACTION&object=TSA-REDUCT-RXN |
| TRSARr | pyr | BW | FW | FW | https://ecocyc.org/ECOLI/NEW-IMAGE?type=REACTION&object=TSA-REDUCT-RXN |
| TRSARr | succ | BW | FW | FW | https://ecocyc.org/ECOLI/NEW-IMAGE?type=REACTION&object=TSA-REDUCT-RXN |
| VALTA | ac | FW | BW | BW, main path for valine synthesis | https://ecocyc.org/ECOLI/NEW-IMAGE?type=PATHWAY&object=VALSYN-PWY |
| VALTA | fru | FW | BW | BW, main path for valine synthesis | https://ecocyc.org/ECOLI/NEW-IMAGE?type=PATHWAY&object=VALSYN-PWY |
| VALTA | glc | FW | BW | BW, main path for valine synthesis | https://ecocyc.org/ECOLI/NEW-IMAGE?type=PATHWAY&object=VALSYN-PWY |

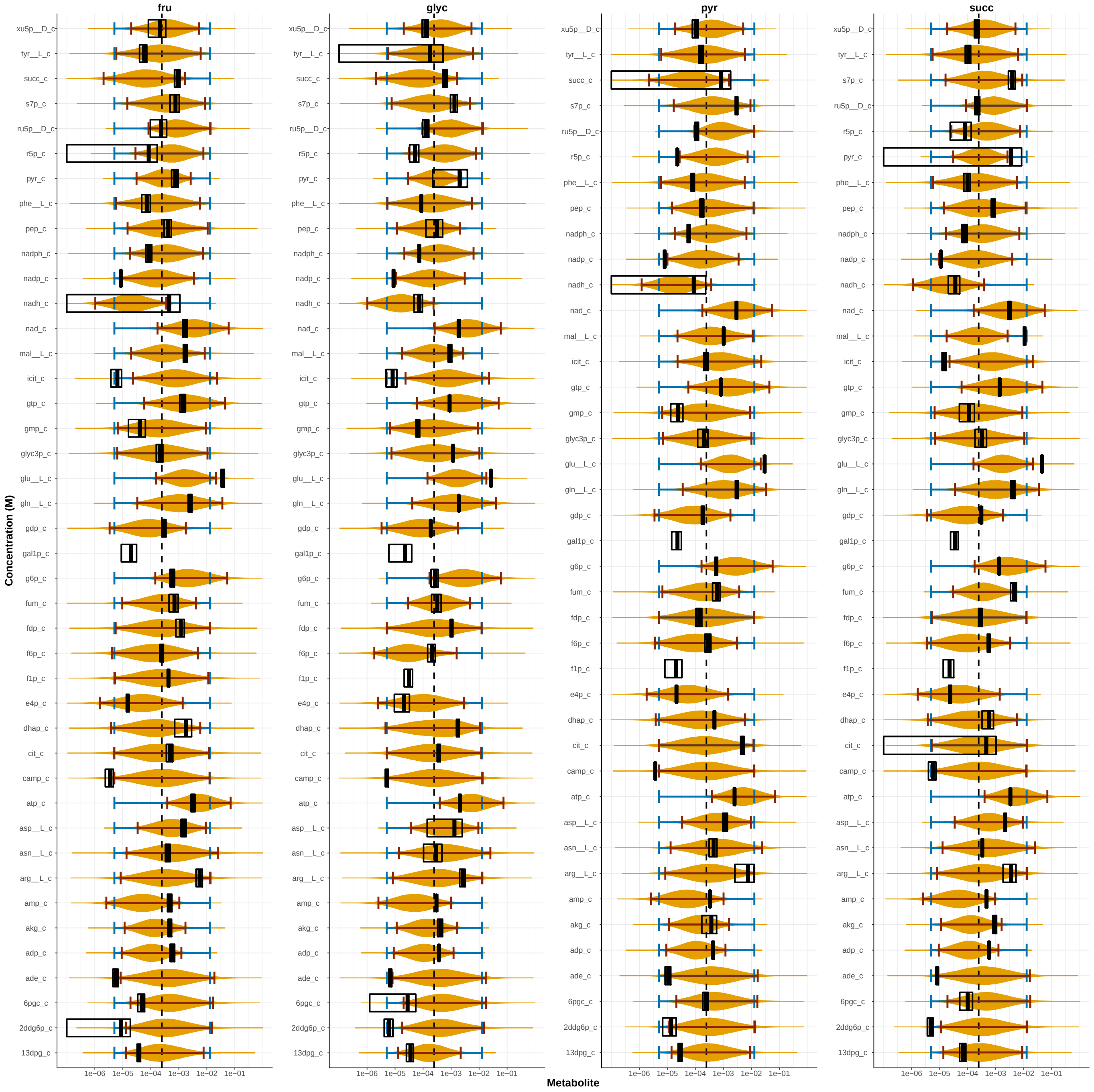


Figure SI 3: Distributions (orange) and 95% confidence intervals (red) of TFS predictions, predicted TMFA ranges (blue) and mean and 95% confidence intervals of the metabolomics data (black) are shown for all metabolites measured in [22]. Some concentrations could not be predicted because all the related reactions were blocked.
